# A Combination of Two Human Monoclonal Antibodies Limits Fetal Damage by Zika Virus in Macaques

**DOI:** 10.1101/2020.01.31.926899

**Authors:** Koen K.A. Van Rompay, Lark L. Coffey, Tania Kapoor, Anna Gazumyan, Rebekah I. Keesler, Andrea Jurado, Avery Peace, Marianna Agudelo, Jennifer Watanabe, Jodie Usachenko, Anil Singapuri, Ramya Immareddy, Amir Ardeshir, Jackson B. Stuart, Stylianos Bournazos, Jeffrey V. Ravetch, Paul J. Balderes, Ivo C. Lorenz, Shannon R. Esswein, Jennifer Keeffe, Pamela J. Bjorkman, Qiao Wang, Charles M. Rice, Margaret R. MacDonald, Michel C. Nussenzweig, Davide F. Robbiani

**Author notes:** Corresponding authors: KKAVR,; MCN,; DFR,.

## Abstract

Human infection by Zika virus (ZIKV) during pregnancy can lead to vertical transmission and fetal aberrations, including microcephaly. Prophylactic administration of antibodies can diminish or prevent ZIKV infection in animal models, but whether passive immunization can protect nonhuman primates and their fetuses during pregnancy has not been determined. Z004 and Z021 are neutralizing monoclonal antibodies to domain III of the envelope (EDIII) of ZIKV. Together the two antibodies protect nonpregnant macaques against infection even after Fc modifications to prevent antibody-dependent enhancement *in vitro* (ADE) and extend their half-lives. Here we report on prophylactic co-administration of the Fc-modified antibodies to pregnant rhesus macaques challenged 3 times with ZIKV during first and second trimester. The two antibodies did not entirely eliminate maternal viremia but limited vertical transmission protecting the fetus from neurologic damage. Thus, maternal passive immunization with two antibodies to EDIII can shield primate fetuses from the harmful effects of ZIKV.

**Significance statement:** Zika virus (ZIKV) infection during pregnancy can cause fetal abnormalities. Vaccines against ZIKV are under development, but because of potential safety concerns due to disease enhancing antibodies, and the time required by active immunization to induce protective antibodies, there is a need to explore alternative strategies. Recombinant monoclonal antibodies can be modified to prevent enhancement of infection, and thus could be an efficacious and safe alternative to vaccines to confer rapid protection. We show that prophylactic administration of two engineered antibodies, Z004 and Z021, to pregnant macaques partially protects against fetal neurologic damage and limits vertical transmission of ZIKV.

---

ZIKV is a mosquito-borne flavivirus that in humans is often asymptomatic but can manifest as febrile illness and rarely as neurologic disease (1, 2). Although ZIKV was discovered in 1947, its devastating effects on fetal health were only recognized during the 2015 outbreak in the Americas (3). ZIKV infection during pregnancy can result in a range of birth defects that predominantly involve the central nervous system including microcephaly (4–6). The rate of adverse outcomes varies but can surpass 40% in some regions (7–10). Similar consequences are observed in macaques, where experimental infection during gestation can lead to fetal neuropathology and demise (11–13).

Although the path to approval is challenging due to the waning epidemic, several ZIKV vaccine candidates have entered clinical trials (14, 15). In preclinical settings, vaccine efficacy was demonstrated in mice and macaques (16, 17), including mouse and macaque pregnancy models (18–21). One of the potential issues with vaccination is that active immunization regimens require time to induce protective immunity and therefore vaccination after conception may be ineffective in preventing fetal disease in the setting of an active outbreak. In addition, there is concern about vaccine-mediated enhancement because features of maternal antibodies that enhance ZIKV infection *in vitro* are associated with an increased risk of human microcephaly and other neurologic defects in humans and macaques (22). Thus, there is a need to develop strategies to limit maternal and fetal pathology that are rapid, effective and safe.

## Results

The combination of human IgG1 monoclonal antibodies Z004 and Z021 significantly reduces plasma viral loads and prevents emergence of viral escape mutations in nonpregnant rhesus macaques challenged with a super-physiologic dose of ZIKV intravenously (10^5^ plaque-forming units, PFU) (23). Similar results were obtained when the antibodies were modified to prevent ADE *in vitro* by mutations that abrogate Fc-g receptor engagement (GRLR; (23, 24)).

To evaluate the protective effect of these antibodies against what is considered to be a more physiologic dose and route of administration, we administered Z004^GRLR^ + Z021^GRLR^ to macaques 24 hours before subcutaneous inoculation with 10^3^ PFU of ZIKV (Fig. 1A) (25, 26). In untreated controls, plasma viremia peaked on day 4-5 after infection, reaching between 10^5^-10^7^ RNA copies/ml (n=4; black in Fig. 1B). In contrast, plasma viremia was undetectable in 3 out of 6 antibody treated macaques, and it was lower and delayed in the remaining 3 animals (green in Figs. 1B-D). Since ZIKV can be detected in macaque tissues even when absent from plasma (13, 27), we measured viral RNA in a panel of 17 tissues (Fig. 2). Viral RNA was evident in multiple tissues in control animals as well as in the 3 animals that showed low and delayed viremia after infection (Fig. 2A, right). In contrast, viral RNA was low or undetectable in tissues from macaques with no detectable plasma viremia (Figs. 2A, left, and B). Thus, prophylactic administration of Z004^GRLR^ + Z021^GRLR^ prevents or reduces viral accumulation in plasma and tissues upon low-dose ZIKV challenge.

**Fig. 1.**
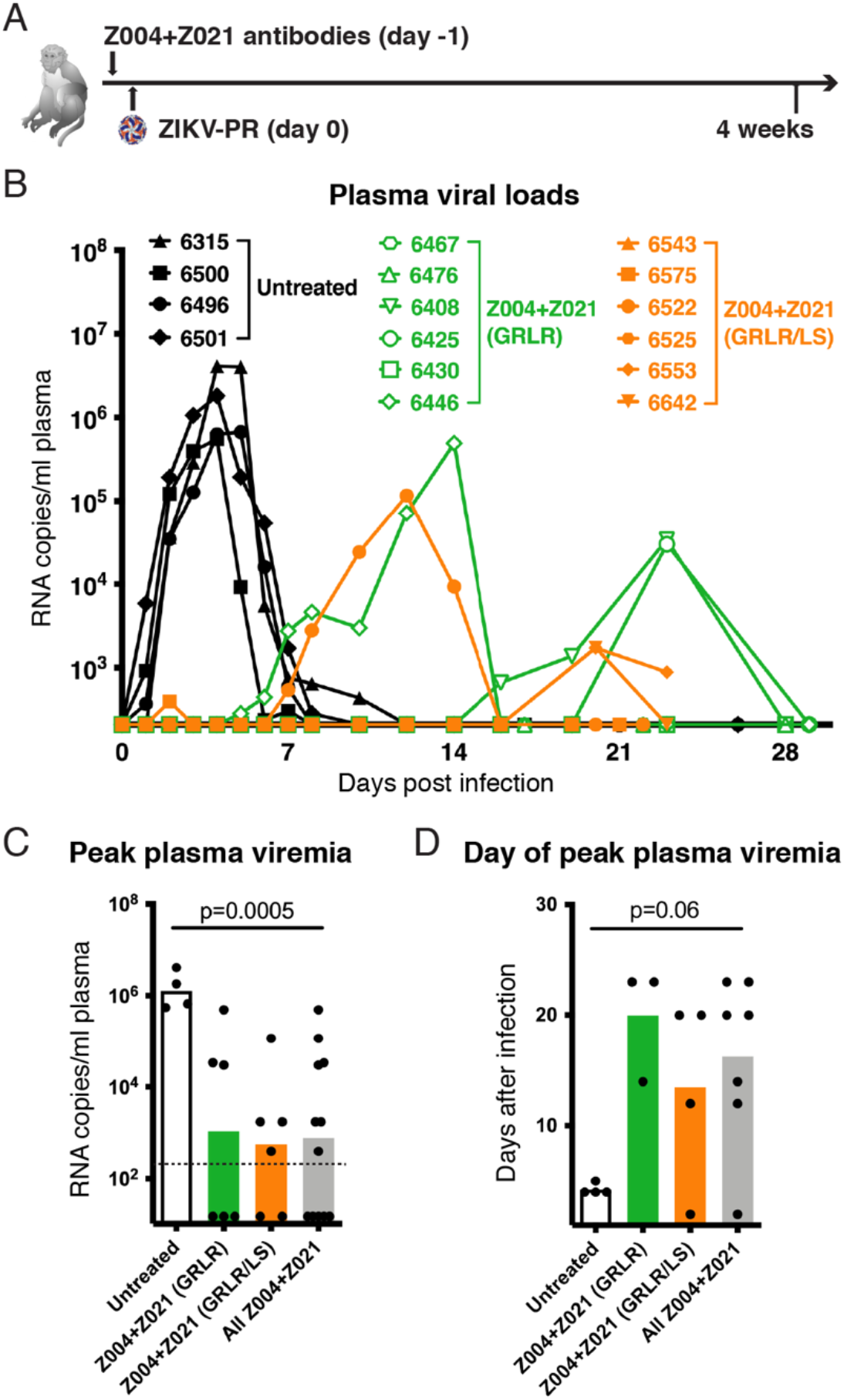
Administration of Z004^GRLR/LS^ + Z021^GRLR/LS^ antibodies reduces magnitude and duration of ZIKV viremia in rhesus macaques. **(A)** Schematic of the experiment. Macaques were administered Fc-modified human monoclonal antibodies Z004 and Z021 one day prior to subcutaneous challenge with 10^3^ PFU of Puerto Rican Zika virus (ZIKV-PR). **(B)** Antibody treatment alters plasma viral loads. Macaques received either the GRLR version of the antibodies (green), the GRLR/LS version (orange), or were left untreated (black). Shown are plasma ZIKV RNA levels over time as determined by qRT-PCR. **(C)** Peak plasma viral RNA levels are decreased in antibody treated macaques (based on panel B). In grey is the peak viral load of both Z004^GRLR^ + Z021^GRLR^ and Z004^GRLR/LS^ + Z021^GRLR/LS^ groups combined. The dotted line represents the limit of detection of the assay and samples with undetectable ZIKV were arbitrarily assigned a value equal to half of the limit of detection. **(D)** Peak plasma viremia is delayed in antibody treated macaques (based on panel B). Bars in C and D represent the mean; the p values were determined with the Mann-Whitney test.

**Fig. 2.**
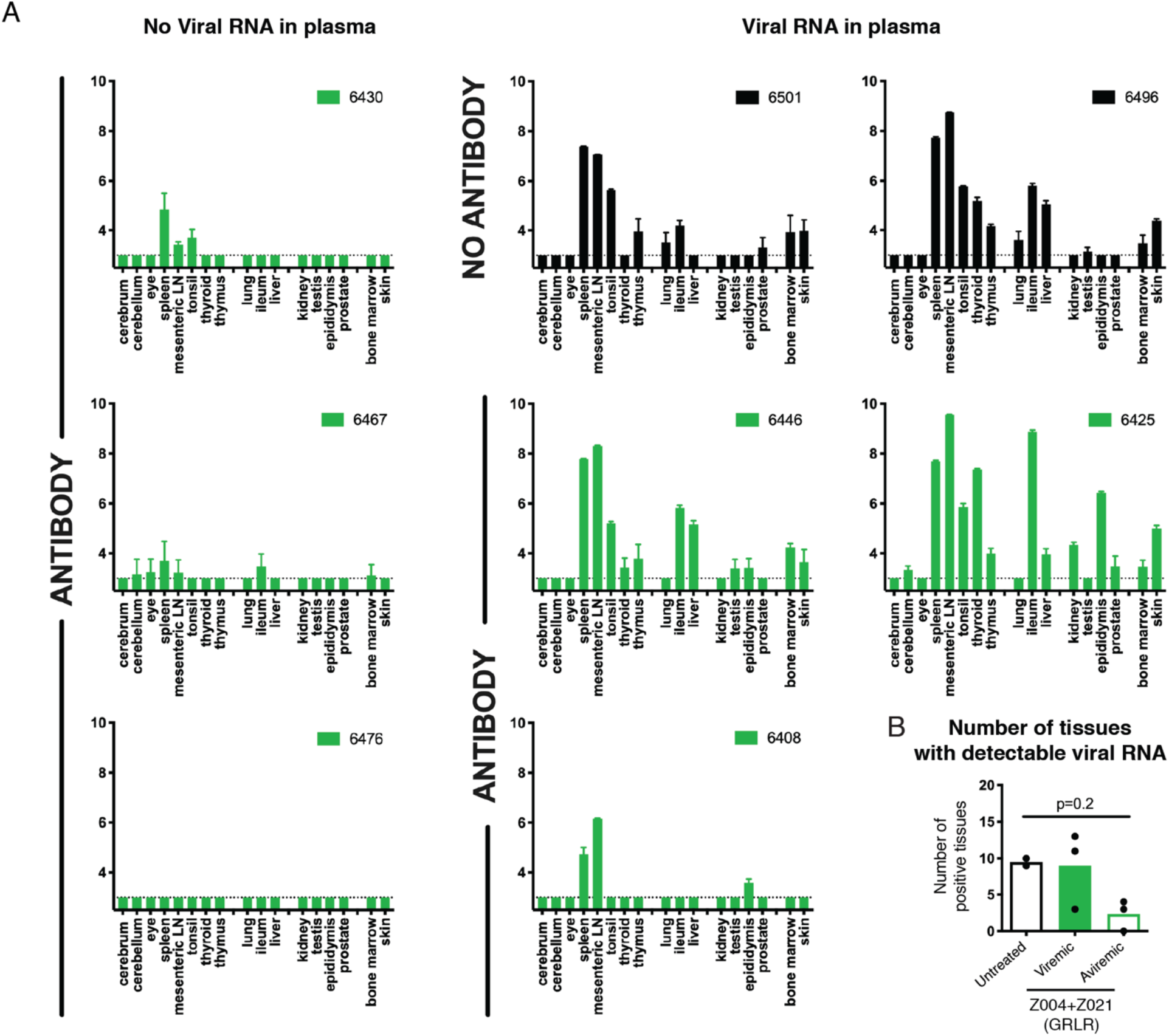
Zika viral RNA levels in tissues are decreased in antibody treated animals. **(A)** Viral RNA was measured in a panel of tissues obtained at euthanasia (day 26-29 post infection). Values represent log_10_ of viral RNA copies per gram of tissue, as determined by qRT-PCR. Data are represented as mean ± SD of triplicate measurements. In green are macaques treated with Z004^GRLR^ + Z021^GRLR^ and in black are untreated controls. **(B)** Shown for each macaque is the number of tissues with detectable viral RNA. Bars represent the mean; the p values was determined with the Mann-Whitney test.

The half-life of Z004 and Z021 can be extended by altering the antibody’s Fc domain (LS mutation; (28, 29)) without altering ZIKV neutralizing activity *in vitro* or *in vivo* (23). To determine whether Z004^GRLR/LS^ + Z021^GRLR/LS^ remain efficacious in macaques, we administered the combination before low-dose subcutaneous challenge with ZIKV. Z004^GRLR/LS^ + Z021^GRLR/LS^ showed extended halflives and their levels in plasma remained more than 4 logs above the *in vitro* inhibitory concentration throughout the monitoring period (Fig. S1). The treatment prevented plasma viremia in 2 of 6 macaques; one macaque had a single episode of low-level viremia (less than 10^3^ viral RNA copies/ml on day 2 post infection), and the remaining 3 had lower and delayed viral loads (orange in Figs. 1B-D) compared to untreated controls. The macaques that developed plasma viremia after administration of Z004^GRLR^ + Z021^GRLR^ or Z004^GRLR/LS^ + Z021^GRLR/LS^ had viruses with mutations in the EDIII region: T309A, T309A/V330L, T309I/V330L and V330L (Figs. 3A-C). To determine whether these substitutions confer resistance to the antibodies, we produced reporter virus particles (RVPs; (30)) with these mutations and evaluated them for sensitivity to the 2 antibodies (Fig. 3D). Both antibodies retained potency against RVPs containing the V330L mutation, which was also found in untreated control animals (Fig. 3A). This mutation is located outside the epitope of either antibody (Fig. 3B). Although T309 is in the epitope of both antibodies (Figs. 3B and C), its mutations affected the potency of Z021 alone. Since T309 mutations did not alter ZIKV sensitivity to the combination of antibodies *in vitro*, they are unlikely to represent virus escape (Fig. 3D). Consistent with these findings, the mutations involved residues other than those altered in the viruses emerging from macaques treated with the individual antibodies (E393 and K394 with Z004, and T335 with Z021; (23) and Fig. S2). We conclude that Z004^GRLR/LS^ + Z021^GRLR/LS^ prevent emergence of viral escape variants found with monotherapy ((23) and Fig. S2) and are as efficacious as their GRLR counterparts against ZIKV in macaques.

**Fig. 3.**
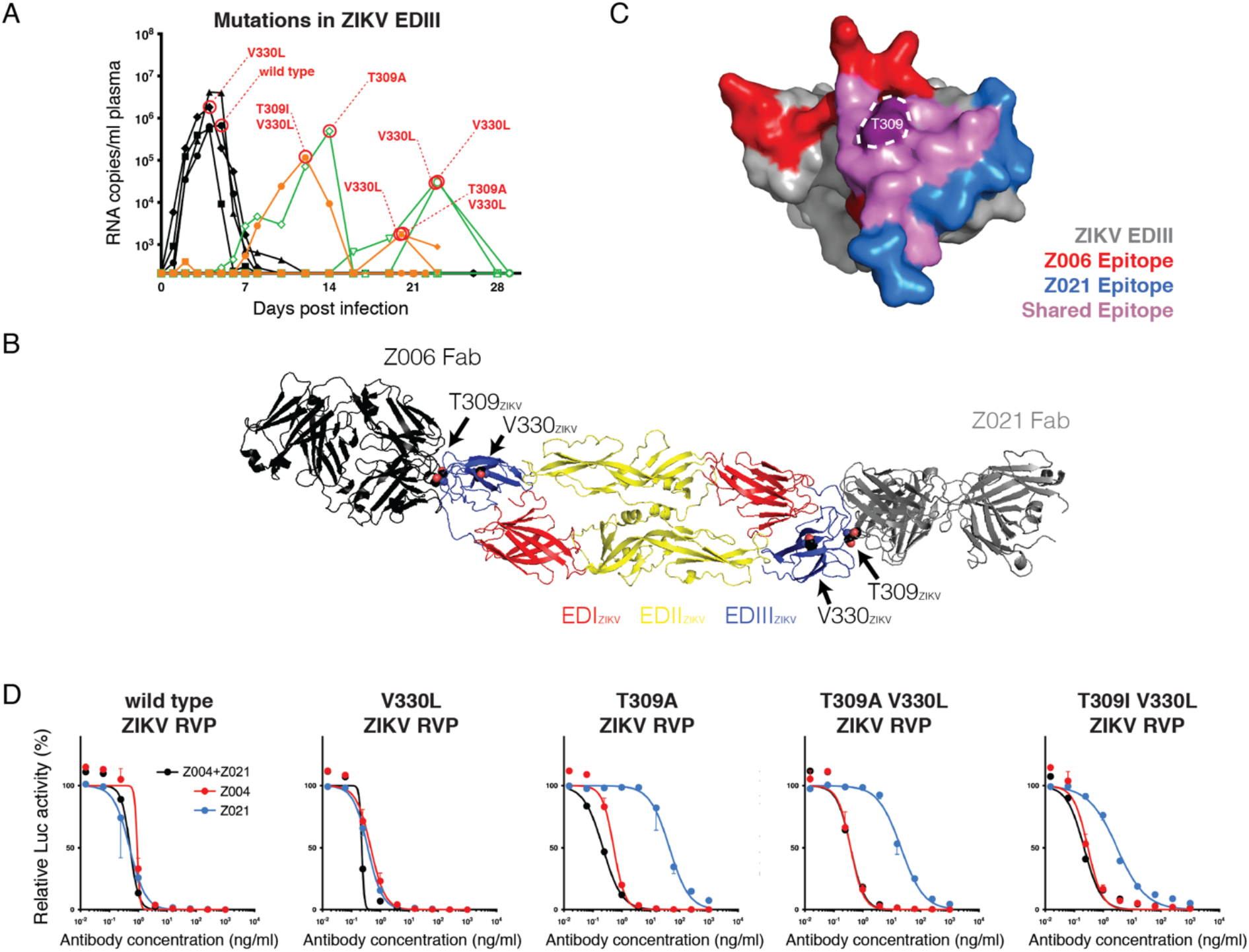
Mutations in Zika virus emerging in plasma from antibody treated macaques remain sensitive to Z004 and Z021. **(A)** Summary of the amino acid changing mutations in the envelope domain III (EDIII) of ZIKV. The virus EDIII region recognized by Z004 and Z021 was amplified and sequenced at peak plasma viremia. Shown is a graphic summary of the identified mutations. Note that V330L was also detected in one untreated macaque. We were unable to amplify the EDIII from one sample with RNA copies/ml below 10^3^. **(B)** The mutations are mapped on the structure of the sE dimer of ZIKV (PDB ID: 5JHM). The structures of the Z004-related antibody Z006 (PDB ID:5VIG) and of Z021 (PDB ID:6DFI) in complex with the EDIII of ZIKV are structurally aligned to the sE dimer to show the location of residues T309 and V330 relative to the binding sites of the antibodies. The ZIKV EDIII from the Z006-EDIII and Z021-EDIII structures are omitted for clarity. **(C)** The epitopes of ZIKV EDIII recognized by the Z004-related antibody Z006 (in red), by Z021 (in blue) and by both antibodies (in purple) are shown. Residue T309 is highlighted. **(D),** Z004, Z021 and the two antibodies together (Z004 + Z021) neutralize RVPs corresponding to ZIKV wild type sequence or ZIKV mutated at the indicated residues. For Z004 + Z021, each of the antibodies is added at the indicated concentration (e.g., 10ng/ml contains 10ng/ml of Z004 plus 10ng/ml of Z021). Data are represented as mean ± SD of triplicates and a representative of two experiments is plotted. Values are relative to isotype control.

ZIKV infection of pregnant macaques is associated with fetal demise and neuropathology (11–13). To evaluate the effect of anti-ZIKV antibodies on fetal development and health in pregnant primates, we performed timed breeding experiments and challenged the dams 3 times with 10^3^ PFU of ZIKV subcutaneously on gestational days (GD) 30, GD 60 and GD 90, which corresponds to 1^st^ and 2^nd^ trimester (normal gestation of rhesus macaques is ~165 days) (21). Macaques in the treatment group received intravenous infusions of Z004^GRLR/LS^ + Z021^GRLR/LS^ antibodies 24 hours before each virus challenge (Fig. 4A). Both dam and fetus were closely monitored; animals were euthanized near the end of gestation for detailed tissue analysis (Tables S1 and S2). During this whole observation period, human IgG plasma levels remained 3 logs or higher than the *in vitro* inhibitory concentration (see Methods and Fig. S3). Untreated macaques develop plasma viremia that peaks between days 2-5 after the first ZIKV inoculation (Fig. 4B, left; (21)). In contrast, out of the 8 antibody-treated dams, 3 had no detectable viremia, while 4 other animals had delayed peak viremia on days 7 (n=1) or 21 (n=3) after the first inoculation and a 5th developed peak viremia on GD90, 30 days after the second ZIKV inoculation (Fig. 4B, right). In addition, viremia was significantly lower and of shorter duration in antibody-treated dams (Figs. 4 B-F). Fetal death (i.e., no heartbeat on ultrasound) was detected in 2 control animals (C03 and C07) on GD 35 and 60 (i.e., 5 and 30 days after the first ZIKV inoculation), with detectable viral RNA in placental and fetal tissues (21). Fetal death was observed on GD51 in one of the antibody-treated macaques (AB02). This occurred one week after undergoing routine amniocentesis and was coincident with a sudden spike of maternal viremia (>10^5^ RNA copies/ml; Fig. 4B) and ZIKV in the amniotic fluid (10^4^ RNA copies/ml) and placental-amniotic tissues (>10^4^ RNA copies/g tissue). In the antibody-treated pregnant dams, mutations in the EDIII region of the emerging viruses were similar to the previously observed mutations that did not alter sensitivity to the antibodies in combination and therefore are unlikely to represent escape (Fig. 3 and Fig. S4). We conclude that the Z004^GRLR/LS^ + Z021^GRLR/LS^ combination prevents or significantly reduces maternal viral loads during pregnancy.

**Fig. 4.**
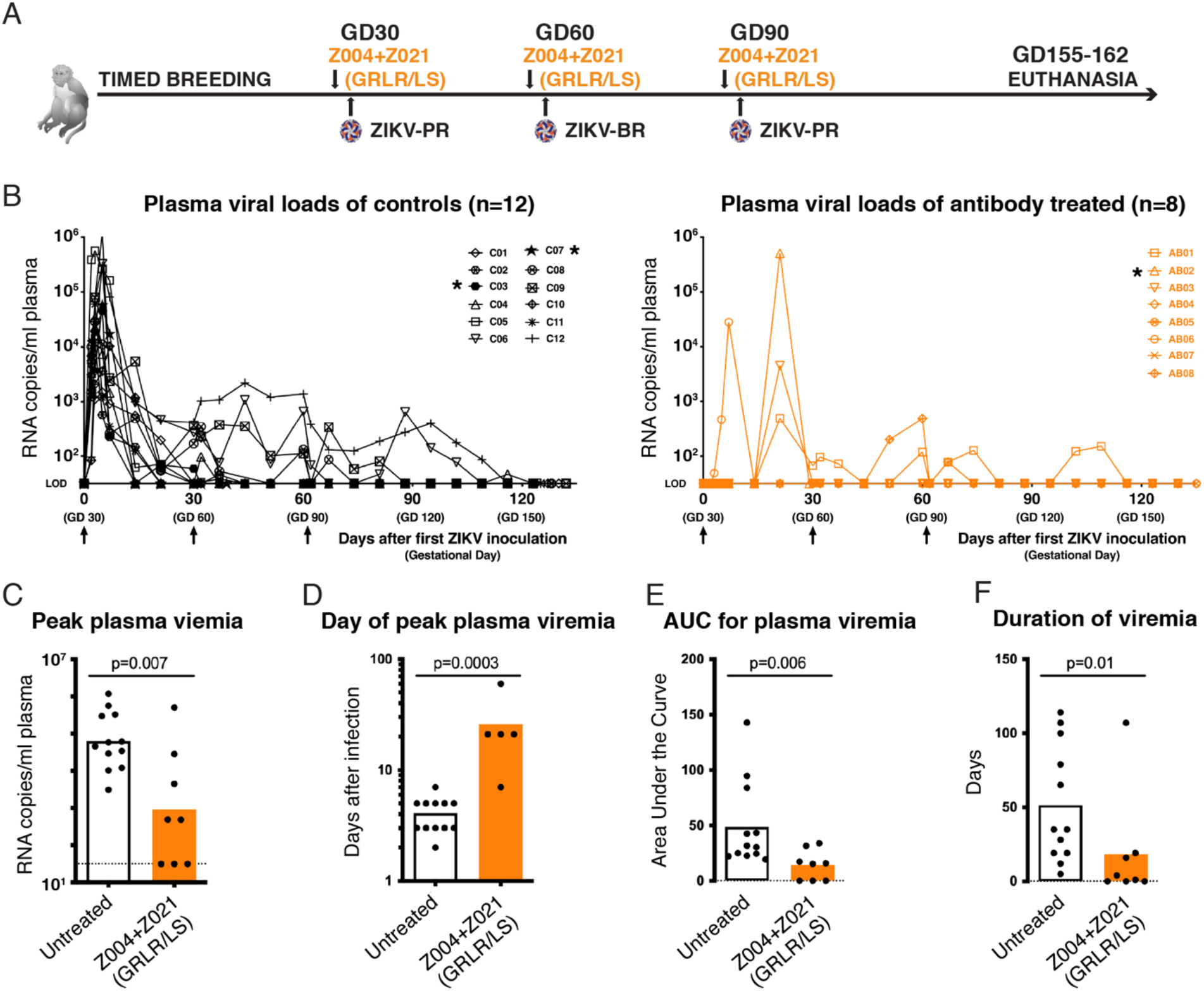
Administration of Z004^GRLR/LS^ + Z021^GRLR/LS^ prevents or reduces ZIKV viral loads in pregnant macaques. **(A)** Schematic of the experiment. Pregnant macaques were administered Z004^GRLR/LS^ + Z021^GRLR/LS^ antibodies one day prior to subcutaneous challenge with 10^3^ PFU of ZIKV on each of gestational days (GD) 30, 60 and 90. PR is Puerto Rican and BR is Brazilian ZIKV. Blood samples were obtained throughout the pregnancy. **(B)** Antibody treatment alters plasma viral loads. Macaques were either untreated (left) or treated with Z004^GRLR/LS^ + Z021^GRLR/LS^ (right). Shown are plasma ZIKV RNA levels over time. LOD is limit of detection. Asterisks indicate animals with early fetal death. **(C)** Peak plasma viral RNA levels are decreased in antibody treated macaques (based on panel b). The dotted line represents the limit of detection of the assay where samples with undetectable ZIKV were arbitrarily assigned a value equal to the limit of detection. **(D)** Peak plasma viremia is delayed in treated animals. **(E)** Overall plasma viral loads are decreased in treated macaques. **(F)** Duration of viremia is reduced in animals that received antibodies. Bars in C-F represent the mean; the p values were determined with the Mann-Whitney test.

Amniocentesis was performed regularly to monitor potential ZIKV transmission to the placental-fetal compartment. Six control animals (50%) had detectable ZIKV RNA in at least 1 amniotic fluid sample. In contrast, except for antibody-treated animal AB02 at time of fetal death, none of the other antibody-treated animals had detectable vRNA in amniotic fluid samples (Fig. S5). Since vertical transmission of ZIKV can cause congenital Zika syndrome, we examined the fetuses in the pregnancies that reached term. Fetal body, placenta, brain and other fetal organ weights were indistinguishable between controls and treatment groups (Fig. S6). In 9 of the 10 control untreated fetuses that made it to the end of gestation, ZIKV RNA was detected in at least one of the 17 examined tissues. These include the maternal-fetal interface, lymphoid, cardio-pulmonary, urinary, gastrointestinal and central nervous systems, and the skin (average of 6 ZIKV RNA positive tissues per fetus; Fig. 5A). In contrast, viral RNA was undetectable in the same fetal tissues from the 7 antibody-treated animals that reached end of gestation (Fig. 5A). Moreover, fetuses that received antibodies had less brain pathology overall, as only 1 out of 7 fetuses (14%) had a brain pathology score of moderate (score of 3) or above, in contrast to 4/10 (40%) of the control fetuses ((21), Figs. 5B, S7 and Table S3). Though its dam had no detectable viremia, it is possible that the moderate brain pathology score in one antibody-treated fetus (AB08-F) reflects earlier infection that was cleared by the time the fetus was harvested. Pathology scores of placenta and amnion were not significantly different between groups (Fig. S7 and Table S3). Finally, there was no correlation between maternal viremia and pathology scores; while aviremic animal AB05 had normal fetus and placenta, macaque AB07 was also aviremic but its fetus and placenta showed mild to moderate pathology scores, respectively; in contrast, the fetus of animal AB06, which had higher than 10^4^ RNA copies/ml at peak maternal viremia, had normal brain findings but moderate placental pathology (Fig. 5B and Fig. S7). We conclude that maternal administration of Z004^GRGR/LS^ + Z021^GRLR/LS^ limits vertical transmission and fetal brain damage by ZIKV.

**Fig. 5.**
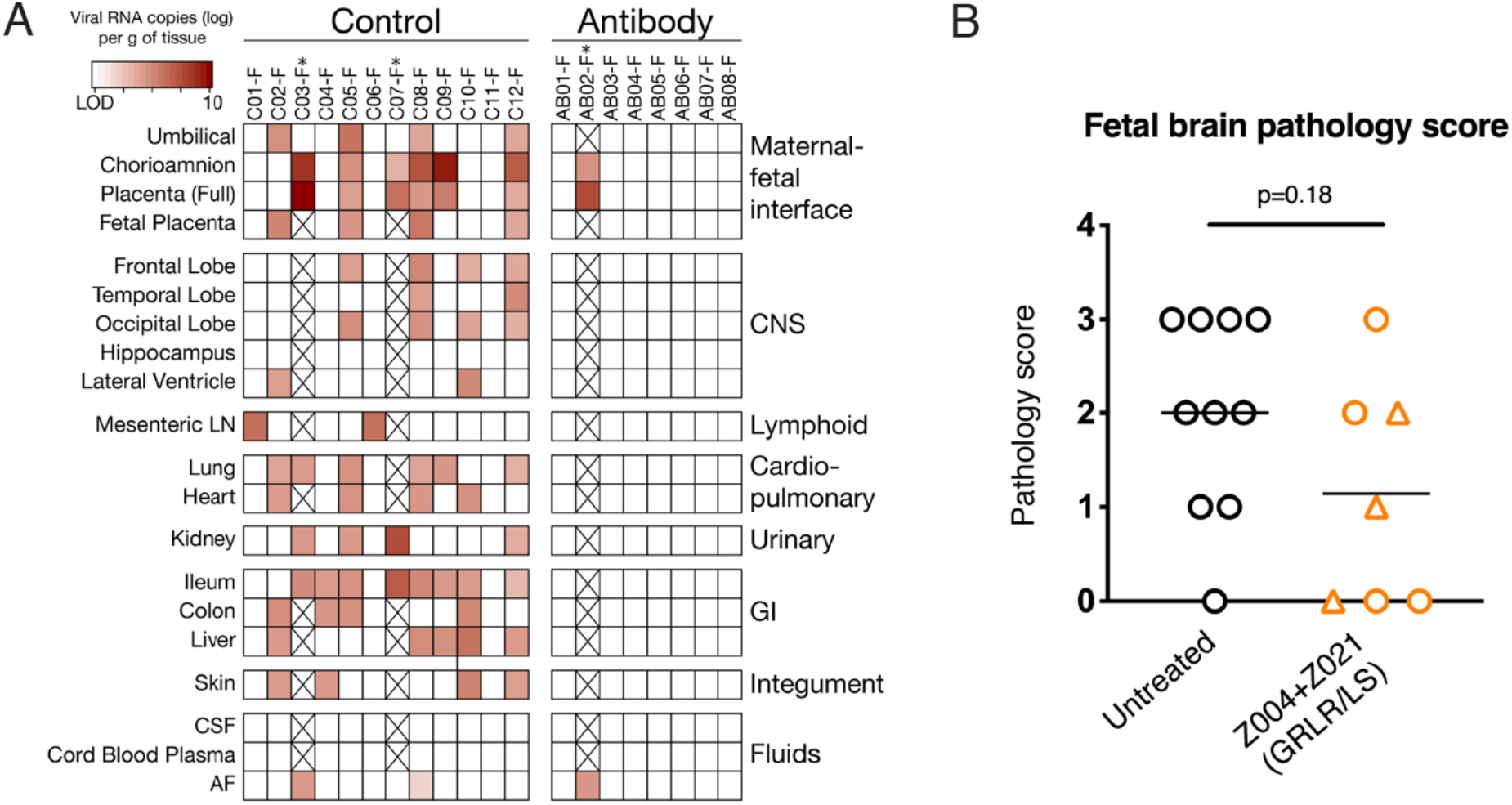
Protection of macaque fetuses by Z004^GRLR/LS^ + Z021^GRLR/LS^. **(A)** With exception of AB02-F (i.e., the fetus of dam AB02), ZIKV RNA is undetectable in placental and fetal tissues from antibody treated dams. Viral RNA was measured in a panel of 17 fetal tissues and 3 fluids harvested at pregnancy termination. LN is lymph node, CSF is cerebrospinal fluid, AF is amniotic fluid, GI is gastro-intestinal, and CNS is central nervous system. * indicates the 2 control animals and one antibody-treated animal with early fetal death. **(B)** Lower fetal brain pathology in animals receiving Z004^GRLR/LS^ + Z021^GRLR/LS^. Triangles indicate fetuses whose dams had no detectable plasma viremia. Horizontal lines represent mean values and the p value was determined with the Mann-Whitney test. Pregnancies that terminated prematurely (GD60 or earlier) were omitted from this panel. Pathology scores of fetal tissues are summarized in Fig. S7.

## Discussion

Recombinant antibodies represent an alternative to vaccines for at-risk populations. Passive immunization reduces or prevents ZIKV infection in nonpregnant rodents and macaques (23, 25, 26, 31, 32). In addition, antibodies diminish ZIKV viral burden in the placenta and fetal organs, and also reduce fetal demise in mouse pregnancy models (33–35). Some of these ZIKV mouse pregnancy models reproduce features of the human disease, including brain pathology (4–6). However, mice are not natural hosts for ZIKV, and the development of pathology typically requires an impaired immune system. As opposed to immunocompetent rodents, macaques can be infected readily with ZIKV and their pregnancies are anatomically and physiologically far more similar to humans (36). Whether antibodies can prevent fetal disease in macaques had not been determined. Our experiments show that the combination of two ZIKV neutralizing human monoclonal antibodies is sufficient to thwart or significantly reduce maternal viremia, prevent virus accumulation in fetal tissues and reduce fetal pathology following repeated ZIKV challenge in macaques. The results have many similarities, but also some differences, with results obtained using the ZIKV DNA vaccine VRC5283 that was tested in the same animal model of congenital ZIKV disease (21).

VRC5283 is a DNA vaccine that produces subviral particles with pre-membrane, membrane and envelope proteins, is immunogenic in phase 1 clinical trial and is currently being evaluated in a phase 2/2b (17, 37, 38). In the macaque pregnancy experiment with similar design to the present study, two doses of vaccine prior to ZIKV inoculation significantly reduced maternal viremia when animals were challenged between 4 days to 1 year after the 2^nd^ immunization (21). Viremia was undetectable in 5 of 13 animals (38%), a rate of protection comparable to the current antibody treatment study (37.5%; n=8). In addition, in the ZIKV DNA vaccine study, the presence or absence of detectable viremia in vaccinated dams correlated with the titer of neutralizing antibody responses at the time of the first ZIKV challenge (i.e., animals with antibody titers above a threshold had no detectable viremia). For vaccinated animals with detectable viremia, the reduced viremia was always early (peak viremia ≤ 5 days after ZIKV inoculation), and short in duration (≤ 14 days after the first ZIKV challenge), consistent with the observed rapid augmentation of antiviral antibody and T-cell mediated immune responses in vaccinated animals after challenge (21).

In the passive immunization studies with Z004 and Z021, antibodies failed to prevent viremia when nonpregnant macaques were inoculated intravenously with a high dose of virus (10^5^ PFU; (23)). In contrast, we achieved an overall 40% protection from detectable viremia in 20 animals infected with a more moderate dose of 10^3^ PFU of ZIKV administered subcutaneously (5 protected out of 12 nonpregnant macaques and 3 protected out of 8 pregnant dams). This dose is still higher than the typical dose received from a mosquito bite, which in laboratory experiments was ~30 PFU for both the ZIKV isolates used in this study (39). This protection was observed irrespective of Fc domain mutations that modify Fc receptor binding. Neutralizing antibody levels on day of challenge were similar between antibody-treated animals with or without detectable viremia, and were ≥ 3 logs higher than the in vitro inhibitory concentration. However, animals showing low levels and even no ZIKV in plasma had detectable viral RNA in tissues. This indicates that in some animals the virus was able to spread despite high levels of the neutralizing antibodies in serum. These results contrast with earlier reports of sterilizing protection by antibodies under similar virus challenge conditions (subcutaneous inoculation with 10^3^ PFU). However, sampling was less frequent and over a shorter time period in those studies and therefore transient or delayed viremia may have gone undetected (25, 26). The delay in viremia in the current study, despite high levels of the 2 neutralizing antibodies in the circulation, suggests that following moderate to high-dose ZIKV inoculation, virus can still reach tissues and initiate replication, either because in some tissues antibodies did not penetrate sufficiently, or because certain cell types became infected via mechanisms (e.g. via different cell receptors) that are insufficiently captured in the standard *in vitro* neutralization assays. The delayed peak viremia (7-60 days post-inoculation) in the pregnant dams likely reflects burst of virus replication in such reservoirs, which then triggered the development of active immune responses. The observation of early fetal losses in 2 control fetuses and one antibody-treated dam, with high levels of virus in fetal or placental tissues, and rapid disappearance of plasma viremia following fetectomy, indicates the fetal-placental compartment as a possible reservoir that is shielded from antiviral immune responses. The observation of delayed viremia also in antibody-treated nonpregnant macaques suggests the existence of additional reservoirs, of which the identification will improve our knowledge of ZIKV pathogenesis and our ability to develop intervention strategies.

Despite these differences in the kinetic of viremia, early versus delayed viremia in vaccinated versus antibody-treated pregnant dams, the reduced maternal viremia and associated improved fetal outcomes were similar in the two studies (21). Thus, the VRC5283 vaccine and the Z004^GRLR/LS^ + Z021^GRLR/LS^ antibodies are comparably efficacious vis-à-vis fetal protection, with each strategy having their intrinsic advantages and limitations in terms of speed of induction, potential for enhancement and durability of protective immunity.

Dengue is a flavivirus closely related to ZIKV. Dengue virus re-infection can lead to antibody-enhanced disease by a mechanism that is associated with serologic cross-reactivity between different dengue serotypes (40–42). Antibodies to dengue also cross-react with ZIKV and experiments in mice and with human placental explants have shown that ZIKV infection can be enhanced by antibodies in these systems (43–46). Conversely, antibodies to ZIKV that are maternally acquired cause more severe disease in mice infected with dengue (47). Finally, features of the antibodies that increase ZIKV infection in *in vitro* ADE assays correlate with an enhanced risk of Zika microcephaly in humans and brain pathology in macaque (22). Thus, even though preliminary epidemiologic studies in humans do not show ADE of ZIKV (48), the potential for enhancement is a safety concern for ZIKV vaccine development that clinical trials need to address. Since monoclonal antibodies that are Fc modified to prevent ADE can shield against ZIKV infection and fetal damage, they represent a potentially safe and efficacious alternative intervention strategy in the face of a ZIKV outbreak.

## Acknowledgments

We thank Kai-Hui Yao and Daniel Yost at Rockefeller University, as well as A. Gibbons, M. Allen, V. Bakula, M. Christensen, I. Cazares, W. von Morgenland, the veterinary staff, pathology staff, and the staff of Colony Management and Research Services, and Clinical Laboratory at the California National Primate Center for expert assistance. This work was supported by awards from NIH U19AI111825, P01AI138938 and UL1TR001866 (to DFR), R01AI037526, UM1AI100663, U19AI111825, P01AI138938 and UL1TR001866 (to MCN), funding by Lyda Hill Philanthropies (to MCN), grants R01AI124690 and U19AI057229 (CCHI pilot project), The Rockefeller University Development Office and anonymous donors (to CMR and MRM), the Office of Research Infrastructure Programs/OD (P51OD011107), start-up funds from the Pathology, Microbiology and Immunology Department (to LLC) and R21AI129479-S (to KKAVR). Support was also provided by the Robertson Therapeutic Development Fund (to DFR and MCN) and a scholarship by the Studienstiftung des Deutschen Volkes (to TK). MCN is an HHMI Investigator.

## Author contributions

KKAVR analyzed and interpreted the data, edited the manuscript, and supervised JW, JU, RI in the laboratory management of the macaque experiments; AA assisted with data analysis/graphing. LLC edited the manuscript, analyzed data and supervised AS and JBS in viral load determination and sequencing of virus escape mutations. TK cloned plasmids to produce mutant ZIKV RVPs, performed RVP neutralization experiments together with QW, analyzed data, and measured human IgG in macaque plasma together with MA. AG produced and purified antibodies for macaque, mouse and *in vitro* experiments. RIK performed necropsies and histopathological evaluations. AJ, AP and SRE produced and evaluated ZIKV RVPs for neutralization by antibodies under the supervision of MRM and CMR. SB and JVR contributed genetically humanized mice and performed related experiments. PB and ICL provided Z004^GRLR/LS^ and Z021^GRLR/LS^ antibodies. JK and PJB performed *in silico* structural analyses. CMR and MRM supervised, interpreted experimental results and edited the manuscript. MCN and DFR supervised, designed and interpreted experiments and wrote the paper.

## Materials and Methods

### Cell lines

Human embryonic kidney HEK-293-6E cells were cultured at 37°C in 8% CO_2_, shaking at 120 rpm. The other cell lines were cultured at 37°C in 5% CO_2_, without shaking: human hepatocytes Huh-7.5 cells were grown in Dulbecco’s Modified Eagle Medium (DMEM) supplemented with 1% nonessential amino acids (NEAA) and 5% FBS; human Lenti-X 293T cells were grown in DMEM supplemented with 10% FBS.

### Antibodies

The human IgG1 antibodies Z004 and Z021 and their Fc-variants (GRLR and GRLR/LS) were previously described (23, 30). Z004^GRLR^ and Z021^GRLR^ were prepared by transient transfection of mammalian HEK-293-6E cells with equal amounts of immunoglobulin heavy and light chain expression vectors. The supernatant was harvested after seven days and the antibodies purified using Protein G Sepharose 4 Fast Flow. LPS was removed with TritonX-114 and the antibodies concentrated in PBS. Z004^GRLR/LS^ (TDI-Y-001) and Z021^GRLR/LS^ (TDI-Y-002) were produced by WuXi and provided by the Tri-Institutional Therapeutics Discovery Institute (TDI, New York). In TDI-Y-002, the Z021 heavy chain variable sequence was modified (C50V).

### Macaque infection experiments

#### Animals and care

All rhesus macaques (*Macaca mulatta*) in the study were born and raised in the conventional (i.e., not specific pathogen free) breeding colony at the California National Primate Research Center (CNPRC; Supplementary Table S4). While none of the animals were positive for type D retrovirus, SIV or simian lymphocyte tropic virus type 1, two animals in the pregnancy control group (C02 and C07) were seropositive for West Nile virus due to prior outdoor housing. All animals in the pregnancy experiment had prior pregnancies (range 2-10).

The CNPRC is accredited by the Association for Assessment and Accreditation of Laboratory Animal Care International (AAALAC). Animal care was performed in compliance with the 2011 *Guide for the Care and Use of Laboratory Animals* provided by the Institute for Laboratory Animal Research. Macaques were housed indoor in stainless steel cages (Lab Product, Inc.) whose sizing was scaled to the size of each animal, as per national standards, and were exposed to a 12-hour light/dark cycle, 64-84°F, and 30-70% room humidity. Animals had free access to water and received commercial chow (high protein diet, Ralston Purina Co.) and fresh produce supplements. The study was approved by the Institutional Animal Care and Use Committee of the University of California, Davis.

##### Experiments with nonpregnant macaques

All 18 rhesus macaques (*Macaca mulatto*) were healthy juvenile males and females (~2 to 4 years of age). When necessary for inoculations or sample collections, macaques were immobilized with 10 mg/kg ketamine hydrochloride (Parke-Davis) injected intramuscularly after overnight fasting. Z004 and Z021 antibodies were administered by the intravenous (i.v.) route to macaques at doses of 15 mg/kg body weight 24 hours before subcutaneous inoculation with Puerto Rican ZIKV (ZIKV-PR; 10^3^ PFU, strain PRVABC-59; GenBank KU501215). For the experiments in Figure S2, two macaques were administered antibody Z021 (with wild type Fc) and infected intravenously with high dose of Brazilian ZIKV (ZIKV-PR; 10^5^ PFU, strain SPH2015; GenBank KU321639.1) similar to previously reported (23). Macaques were evaluated at least daily for clinical signs of disease including poor appetence, stool quality, dehydration, diarrhea, and inactivity. None of the animals that received the antibodies developed any clinical signs, suggesting a good safety profile. Animals were sedated for antibody administration (minus 24h), at time zero (virus inoculation), daily for 7 to 8 days, and then every few days for sample collection. EDTA-anti-coagulated blood samples were collected using venipuncture and plasma isolated. At the end of the study, animals were euthanized for tissue collection. The samples were processed, and the viral RNA measured from plasma and tissues as described below for pregnant macaques.

##### Experiments with pregnant macaques

###### Time-mated breeding and pregnancy selection

For time-mated breeding, the female macaques were monitored for reproductive cycle and at the time of optimal receptiveness, temporarily housed with breeding males to induce pregnancy. Gestational ages were determined from the menstrual cycle of the dam and the fetus length at initial ultrasound compared to growth data in the CNPRC rhesus macaque colony. Fetal health and viability were rechecked via ultrasound immediately before the first ZIKV inoculation (~ GD 30) and regularly thereafter.

###### Antibody treatment and virus inoculations

Z004 and Z021 antibodies were administered to macaques by the i.v. route at doses of 15 mg/kg body weight each by 24 hours before each virus inoculation. Each pregnant animal was inoculated with virus 3 times. Whereas the normal gestation of rhesus macaques is 165 days, inoculations occurred at approximately GD 30, 60 and 90, corresponding to first and second trimester of human gestation. The GD 30 and GD 90 inoculations were done with a 2015 Puerto Rico isolate (PRVABC-59; GenBank KU501215), while the GD 60 inoculation was done with a 2015 Brazil isolate (strain Zika virus/H.sapiens-tc/BRA/2015/Brazil_SPH2015; GenBank KU321639.1), the same strain as tested earlier in pregnant and nonpregnant animals (11, 27). The use of two strains was intended to mimic an endemic area where different variants may circulate. Aliquots of both virus stocks were kept frozen in liquid nitrogen, and a new vial was thawed shortly before each inoculation. For each inoculation, the inoculum was adjusted to 1,000 PFU in 0.5 ml of RPMI-1640 medium, then kept on wet ice and injected subcutaneously to simulate the route of mosquito feeding. This dose is higher than the typical dose received from a mosquito bite, which in laboratory vector competence experiments with Californian *Aedes aegypti* was ~30 PFU for both the ZIKV isolates used in this study (39).

###### Sample collection and clinical observations and monitoring

Macaques were evaluated twice daily for clinical signs of disease including poor appetence, stool quality, dehydration, diarrhea, and inactivity. When necessary, macaques were immobilized with ketamine hydrochloride (Parke-Davis) at 10 mg/kg and injected intramuscularly after overnight fasting. Animals in both the ZIKV treated and placebo cohorts were sedated at days 0 (time of first virus inoculation; approximately GD 30), 2, 3, 5, 7, 14, 21, 30 (2^nd^ ZIKV inoculation at GD60), 32, 37, 44, 51, 60 (3^rd^ ZIKV inoculation; GD90), 62, 67, and then weekly until time of euthanasia between GD155-162, for sample collection and ultrasound monitoring of fetal health. The antibody-treated animals had additional time points of sedation for blood collection and IV antibody administration one day before each ZIKV inoculation. EDTA-anti-coagulated blood was collected using venipuncture at every time point for complete blood counts (with differential count), and a separate aliquot of blood was centrifuged for 10 min at 800 g to separate plasma from cells. The plasma was spun an additional 10 min at 800 g to further remove cells, and aliquots were immediately frozen at −80°C.

Ultrasound guided amniocentesis was conducted starting at day 14 after inoculation (GD44), and then at all time points listed above with exception of days 32 and 62 after initial infection. The amniocentesis was conducted using sterile techniques by inserting a 22 gauge, 1.5 inch spinal needle into the amniotic sac. The fetal heart rate was obtained before and after amniocentesis. The area of umbilical entry through the amniotic sac was always avoided to prevent damage to the umbilical arteries and vein. Whenever possible, placental tissue was avoided during the collection of amniotic fluid. Amniotic fluid was spun to remove cellular debris, and the supernatant was aliquoted and immediately cryopreserved at −80°C for viral RNA assays.

###### Necropsy and tissue collection

All necropsies were performed by a board-certified anatomic pathologist and a pathology technician. For animals that had early fetal loss, the procedures described below were performed to the best extent possible based on fetal size. Hysterotomy was performed by a veterinary surgeon on the pregnant macaques under inhalation anesthesia. After collection of amniotic fluid and cord blood, the fetus was euthanized with an overdose of sodium pentobarbital (≥120 mg/kg). Fetal and organ weights were measured with a scale, and crown-rump length, biparietal diameter, head height, head length, and femur length, were derived from caliper measurements. A detailed tissue dissection was performed. Except for some animals that had early fetal loss and were maintained several weeks after removal of the fetus, all other mothers were euthanized shortly after their fetus with an overdose of sodium pentobarbital for tissue collection.

Each tissue was grossly evaluated *in situ*, and then excised, with further dissection with separate forceps and razor blades for each tissue to minimize risks for cross-contamination. Tissues were collected for viral analyses in RNA*later* (according to manufacturer’s instructions); extra available samples were snapfrozen and stored at −70°C. Tissues were also preserved in 10% neutral buffered formalin and routinely paraffin-embedded and slides were created and routinely stained with hematoxylin and eosin (H&E).

###### Isolation and quantitation of viral RNA from fluids and tissues for determination of infection status

ZIKV RNA was isolated from samples and measured in triplicate by qRT-PCR according to methods described previously (27) modified to increase the initial volume of sample tested from 140 to 300 μl to increase sensitivity. According to the volume available, the limit of detection for plasma and amniotic fluid ranged from 1.5 to 2.3 log_10_ viral RNA copies per ml fluid. RNAlater preserved tissues were homogenized to a liquid state with glass beads (Fisher Scientific) or a 5 mm steel ball (Qiagen). For tissues, the limit of detection (LOD) varied depending on the weight of tissue sampled with a mean of 3.5 log_10_ RNA copies/g of tissue, determined according to the calculations described in (49).

For maternal plasma, amniotic fluid, fetal cord blood and fetal cerebrospinal fluid, all samples from both control and vaccine group animals that could be collected were tested. For the maternal tissue samples, as maternal infection was not the main focus of this study, a limited selection of lymphoid tissues most likely to have viral RNA based on prior studies was tested (11, 27), with addition of uterus due to proximity to the placenta and fetus. For the analysis of fetal tissue samples for viral RNA, we started our analysis with the control fetuses and tested 26 fetal and maternal-fetal-interface (MFI) tissues, consisting of tissues most likely to contain viral RNA based on our prior study (11). Next we generated a heatmap and selected a list of 17 tissues consisting of those most likely to be infected and diagnosing the most infections in the control fetuses, with addition of brain regions most likely to have histological lesions in the current study (such as hippocampus), or found to be infected in human fetuses affected by CZS (50). The criteria to define infection were used systematically for both control and antibody treated group fetuses: a fetus was considered ZIKV-infected if at least one tissue had a consistent qRT-PCR signal (3/3 replicates positive for viral RNA); this situation applied to 11 of the 12 control fetuses. In a rare case, when a fetus only had a sample with inconsistent qRT-PCR signal (1 out of 3 replicates positive) while all other tissues were negative, retesting was performed, generally on a different aliquot. If the retest result was negative (3/3 replicates), the sample was considered not to contain detectable ZIKV RNA, if this occurred for all tissues from a fetus, the fetus was considered not infected. One control fetus (C07-F) and one antibody group fetus (AB05-F) met these criteria and were therefore considered uninfected.

###### Histopathology

Sections of fixed and embedded fetal tissues were stained with hematoxylin and eosin (H&E) and evaluated by a board-certified anatomic pathologist. Scoring of critical fetal tissues was done according to criteria outlined in Table S3.

### Virus sequencing

For the detection of virus mutations at the envelope domain III (EDIII) region, ZIKV RNA was extracted at peak plasma viremia and Qiagen One-Step RT-PCR was performed using either of two primer sets targeting sequences surrounding the ZIKV EDIII region, as previously detailed (23). Due to low levels of viremia in some samples, reliable EDIII sequence information was not always obtained in samples with amounts below 10^3^ RNA copies/ml plasma. Mutations reported are based on changes in the majority nucleotide in sequencing chromatograms; concordant sequences in both directions were used to identify mutations.

### Reporter virus particles (RVPs)

Luciferase encoding RVPs bearing the wild type or mutant ZIKV E proteins of interest were produced by co-transfection of Lenti-X-293T cells with plasmid pWNVII-Rep-REN-IB (WNV replicon expression construct; (51)) in combination with plasmids expressing flavivirus CprME (1 μg of pWNVII-Rep-REN-IB and 3 μg of the CprME plasmid) as previously detailed (30).

The plasmids for expression of wild type ZIKV CprME with E protein sequence corresponding to Puerto Rican ZIKV (strain PRVABC59) that was used in this study was generated from plasmid pZIKV/HPF/CprM*E* (30) by replacing a BspHI/SacII fragment with sequences corresponding to the PRVABC59 strain that were generated by PCR using oligos RU-O-24379 and RU-O-24380 and PRVABC59 derived cDNA as template. The resulting plasmid was designated pZIKV/HPF/CprM*PRVABC59E*. Assembly PCR-based site-directed mutagenesis was then used to introduce in pZIKV/HPF/CprM*PRVABC59E* the V330L, T309A, V330L/T309A and V330L/T309I mutations that resulted in plasmids pZIKV/HPF/CprM*PRVABC59E* (V330L), pZIKV/HPF/ CprM*PRVABC59E* (T309A), pZIKV/HPF/CprM*PRVABC59E* (V330L/T309A) and pZIKV/HPF/ CprM*PRVABC59E* (V330L/T309I). Similarly, the ZIKV RVPs encoding for the T335 mutations (Fig. S2) were generated by modifying plasmid pZIKV/HPF/CprM*E* (30) to introduce the T335A and T335I modifications resulting in pZIKV/HPF/CprM*E* (T335A) and pZIKV/HPF/CprM*E* (T335I), respectively. The primers used for assembly PCR and mutagenesis are listed in Table S5. All PCR- erived plasmid regions were verified by sequencing.

### In vitro neutralization assay

RVP neutralization of luciferase-encoding RVPs by antibodies was performed with Huh-7.5 cells in 96-well plates, in triplicate, as previously detailed (30), and the luciferase activity was read using a FLUOstar Omega luminometer (BMG LabTech). The neutralization capacity of the antibodies against wild type or mutant RVPs was determined by the percentage of luciferase activity obtained relative to RVPs incubated with isotype control antibody 10-1074 (52). Representatives of at least two independent experiments are shown.

### Measurements of human IgG in macaque plasma

ELISA was used for the detection of human IgG as previously detailed (30). Briefly, the biotinylated antihuman IgG capture antibody was added to neutravidin-coated 96-well plates. Upon washes, the plates were blocked with 2% BSA, blotted, and then serial dilutions of the macaque plasma were added to the wells (5 steps of 1:4 dilutions in PBS-T, starting with 1:10). Each plate included serial dilutions of the standard, in duplicates (Z004 IgG, 11 steps of 1:3 dilutions in PBS-T, starting with 10 μg/ml). Plates were incubated for 1 hour at room temperature and washed prior to adding the detection reagent anti-human IgG-HRP. After washes, the reaction was developed with ABTS substrate (Life Technologies).

### Measurements of human IgG in mouse serum

ELISA was used for the detection of human IgG as previously detailed (53). Briefly, FcRn/FcγR-humanized mice were injected i.v. with 100 μg of Z004 or Z021 Fc variants. High-binding 96-well microtiter plates (Nunc) were coated overnight at 4°C with Neutravidin. All sequential steps were performed at room temperature. Plates were blocked for 1 hour with PBS/2% BSA and incubated with biotinylated goat anti–human IgG antibodies for 1 hour (5 μg/mL; catalog 109-066-170, Jackson ImmunoResearch). Serum samples were serially diluted and incubated for 1 hour, followed by incubation with horseradish peroxidase–conjugated anti–human IgG. After washes, the reaction was developed with the TMB peroxidase substrate system (KPL) and stopped with 2 M phosphoric acid. Absorbance at 450 nm was immediately recorded using a SpectraMax Plus spectrophotometer (Molecular Devices), background absorbance from negative control samples was subtracted, and duplicate wells were averaged.

### Statistical analyses

Statistical analyses were with Prism 8 (GraphPad).

