## Supplemental for "A Combination of Two Human Monoclonal Antibodies Limits Fetal Damage by Zika Virus in Macaques"

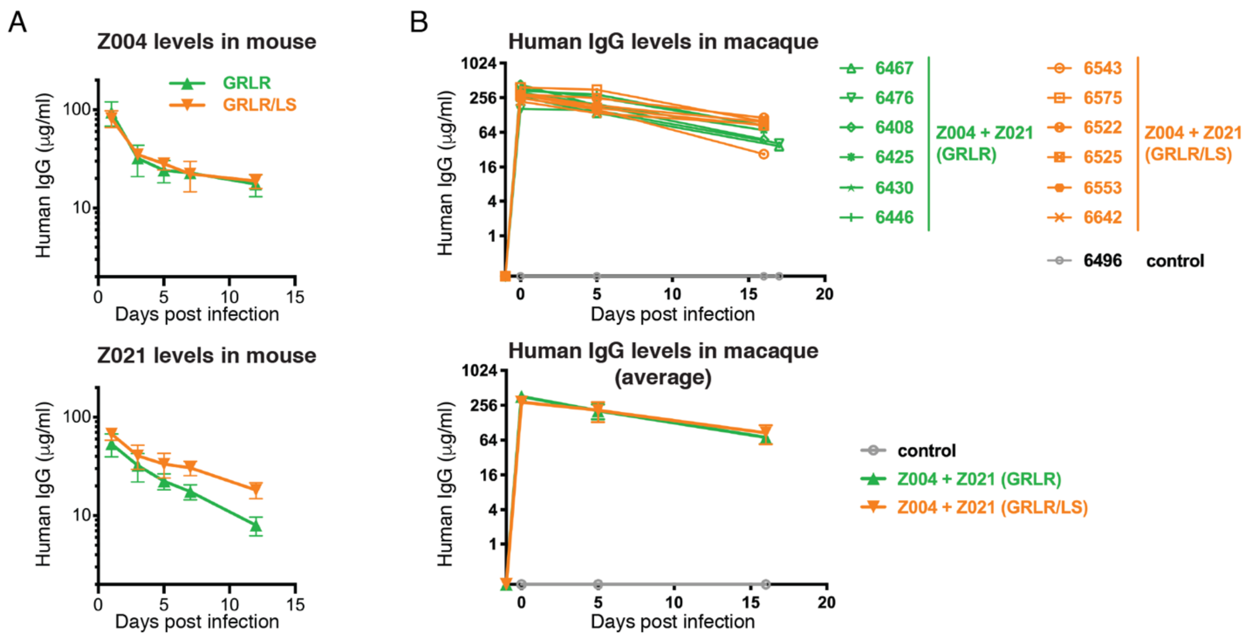


**Fig. S1. Levels of Z004 and Z021 human antibodies in mice and macaques. (A)** Amount of human IgG antibodies in serum over time upon injection of 100µg of either Z004 or Z021 into FcγR/FcRn humanized mice. Four mice per group; shown is the mean ± SD. **(B)** Levels of human IgG antibodies in macaque plasma. The top panel displays human IgGs in individual macaques; the bottom panel shows the mean for each group. Macaques were administered 15 mg/kg of each of the antibodies on day -1. The mean peak antibody levels on the day of infection (day 0) were 364 µg/ml in the $Z004^{GRLR}$ + $Z021^{GRLR}$ group and 289 µg/ml in the $Z004^{GRLR/LS}$ + $Z021^{GRLR/LS}$ group. The antibody levels on day 16 were 71 µg/ml in the $Z004^{GRLR}$ + $Z021^{GRLR}$ group and 85 µg/ml in the $Z004^{GRLR/LS}$ + $Z021^{GRLR/LS}$ group, resulting in plasma half-lives of 6.8 and 9 days, respectively. Samples from macaques 6467 and 6476 (both treated with GRLR antibodies) were excluded from the calculation of the half-life since they were sampled on a different day. Control is plasma from a ZIKV infected, untreated animal.

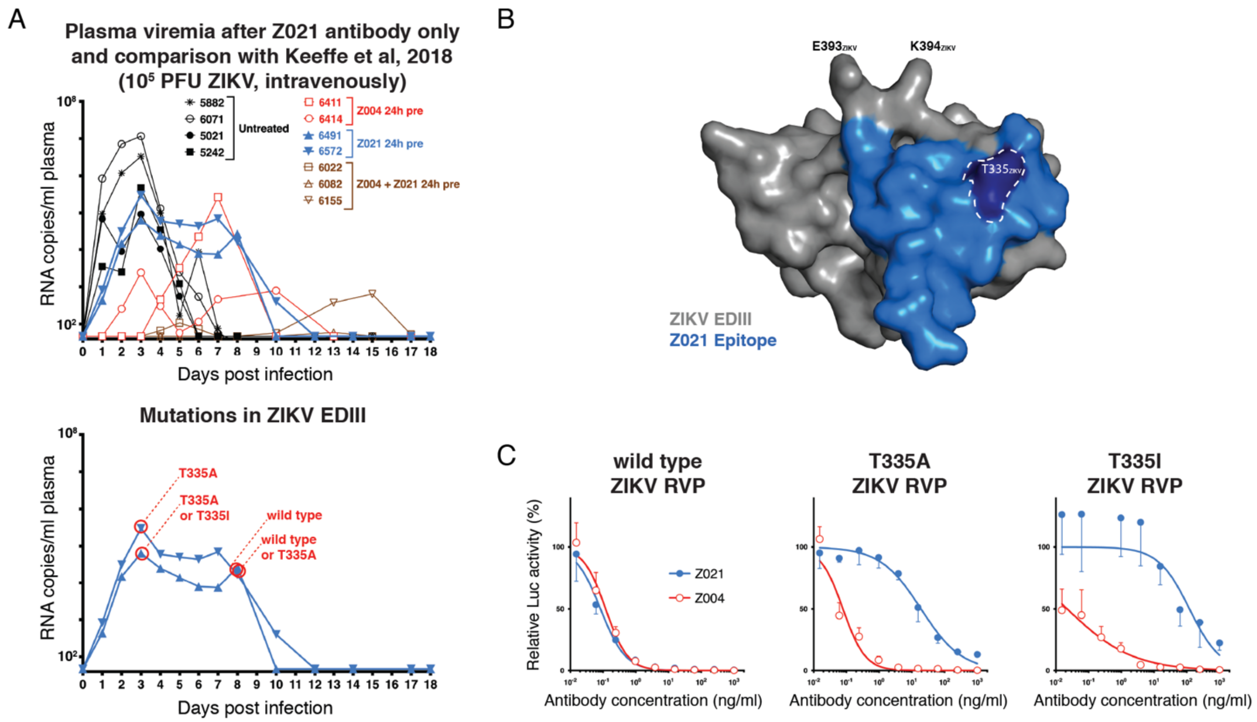


**Fig. S2. Administration of Z021 alone induces the emergence of resistant ZIKV in macaque. (A)** Macaques were administered the Z021 monoclonal antibody 24 hours before intravenous inoculation with $10^5$ PFU of Brazilian ZIKV. The top graph shows the measurement of plasma viremia over time upon Z021 treatment (blue), alongside the values previously reported for Z004 alone, Z004 + Z021, and untreated controls, for which the same virus, dose and route of administration were used (23). The bottom graph shows the time points analyzed that revealed T335 mutations in the virus EDIII region. **(B)** The epitope of ZIKV EDIII recognized by the Z021 antibody is shown in blue (PDB: 6DFI). The T335 residue is highlighted, as well as residues E393 and K394 that when mutated can confer resistance to Z004 (23). **(C)** Z004 similarly neutralizes RVPs corresponding to wild type ZIKV sequence or ZIKV mutated at the T335 residue, while Z021 is less potent against viruses containing T335A or T335I. Data are represented as mean ± SD of triplicates and are relative to isotype control.

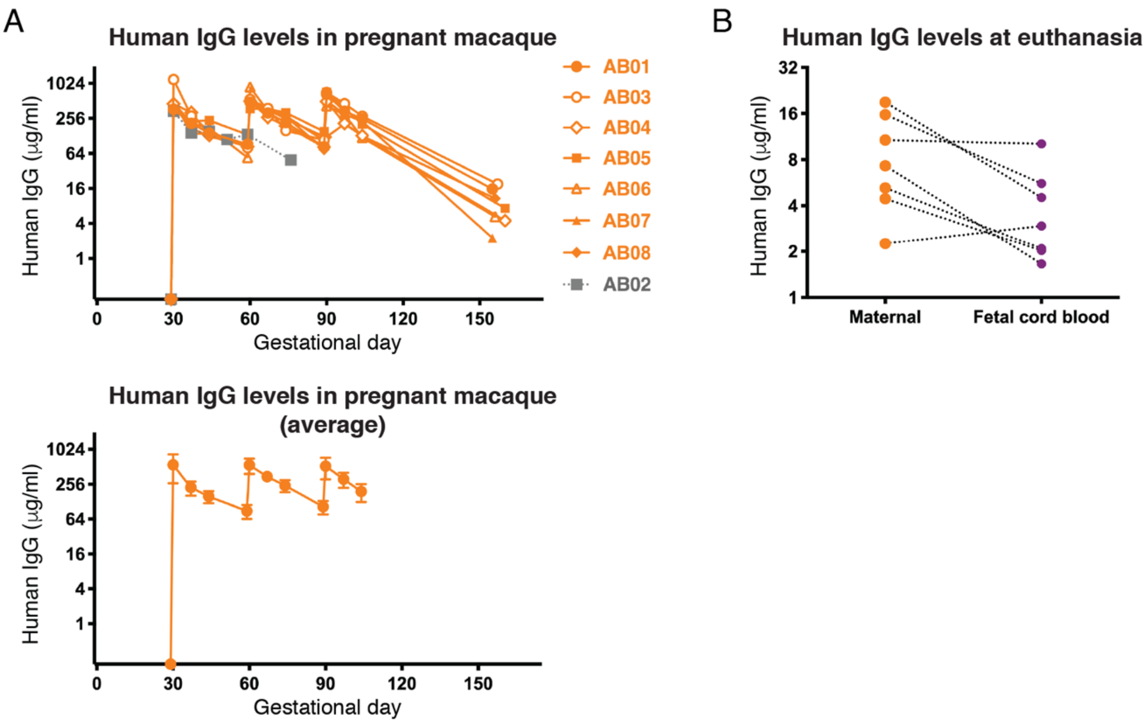


**Fig. S3. Plasma antibody levels of Z004$^{GRLR/LS}$ + Z021$^{GRLR/LS}$ in dams and fetuses. (A)** Levels of human IgG antibodies in plasma of pregnant macaques. The top panel displays human IgGs in individual macaques; the bottom panel shows the mean with standard deviation. Macaques were administered 15mg/kg of each of the antibodies 24 hours before each of the virus challenges, which occurred on GD30, GD60 and GD90. The mean peak antibody levels were 526 µg/ml (GD30), 547 µg/ml (GD60) and 522 µg/ml (GD90); and the calculated half-lives were 11.7 days (GD30-GD59), 12 days (GD60-GD89) and 9.6 days (GD90-GD104). In grey in the top panel are the values for the pregnancy that was prematurely terminated due to fetal loss, which did not receive a 2$^{nd}$ dose of antibodies, and which was excluded from the calculations for the average (bottom panel) and for the half-life values. **(B)** Human IgG levels in the dam and respective fetus (cord blood) at the time of euthanasia.

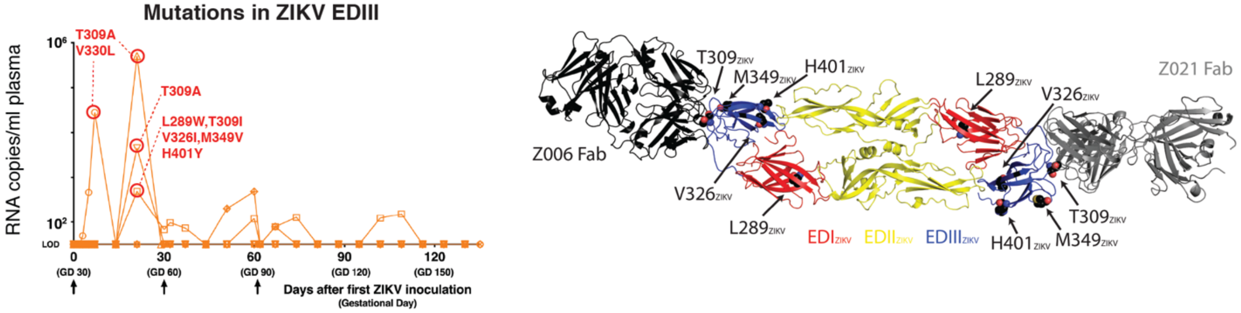


**Fig. S4. Mutations in Zika virus emerge in plasma from antibody-treated pregnant macaques.** On the left is a graphic summary of the identified mutations. Most of the mutations had already emerged in the antibody treated nonpregnant macaques (see Fig. 3). One exception was a virus detected in one animal with mutations resulting in 5 amino acid substitutions (L289W, T309I, V326I, M349V, H401Y); the effect of these mutations on antibody sensitivity could not be evaluated because we were unable to produce RVPs corresponding to these changes. We were unable to amplify the EDIII from most samples with RNA copies/ml below $10^3$. On the right, the mutations are mapped on the structure of the sE dimer of ZIKV (PDB ID: 5JHM). The structures of the Z004-related antibody Z006 (PDB ID:5VIG) and of Z021 (PDB ID:6DFI) in complex with the EDIII of ZIKV are structurally aligned to the sE dimer to show the location of the mutated residues relative to the binding sites of the antibodies. Except for T309, all mutations are outside of the epitopes of both antibodies. The ZIKV EDIII from the Z006-EDIII and Z021-EDIII structures are omitted for clarity.

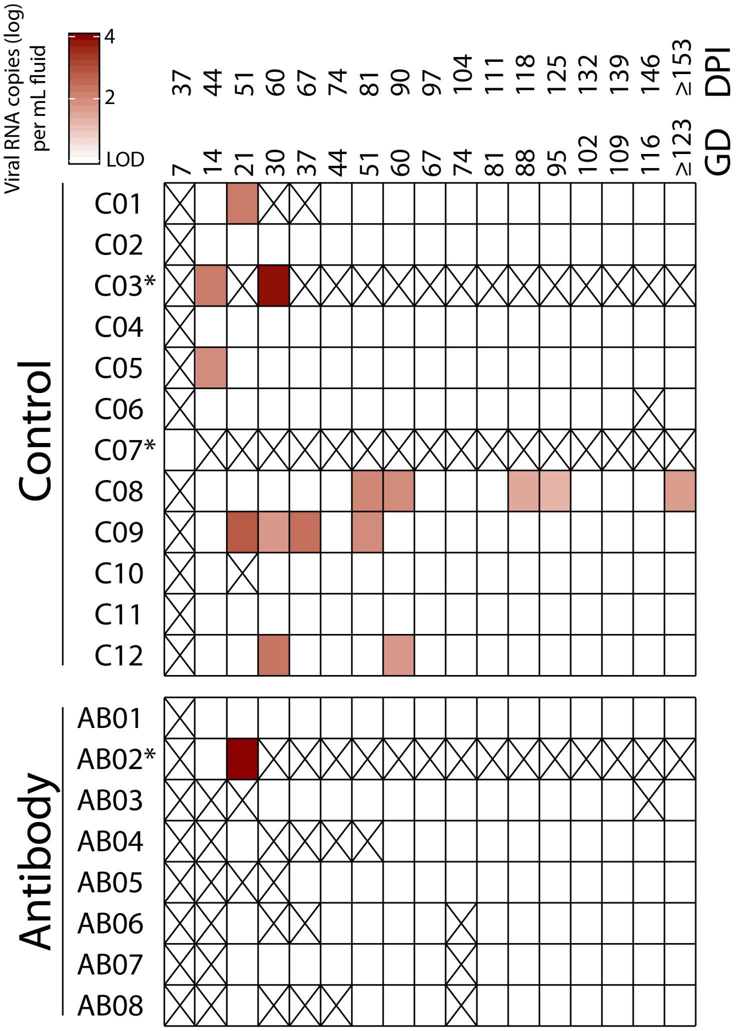


**Fig. S5. Detection of ZIKV RNA in amniotic fluid.** Amniotic fluid was collected regularly by amniocentesis and tested for ZIKV RNA by qRT-PCR. Results are indicated as $\log_{10}$ ZIKV RNA copies/ml amniotic fluid, the mean of triplicate qRT-PCR replicates. Blank cells indicate samples that tested negative, i.e., below the limit of detection (~1.5 to 2.3 $\log_{10}$ ZIKV RNA copies/ml amniotic fluid, depending on volume). The intensity of red highlights the quantity of viral RNA detected. Cross-out indicates samples not tested, because either not collected (GD 37), not able to collect despite amniocentesis attempt, or early pregnancy termination; * indicates the 2 control animals and one antibody-treated animal with early fetal death. The ≥ GD 153 time point is the time of euthanasia. DPI: days post-infection; GD: gestational day.

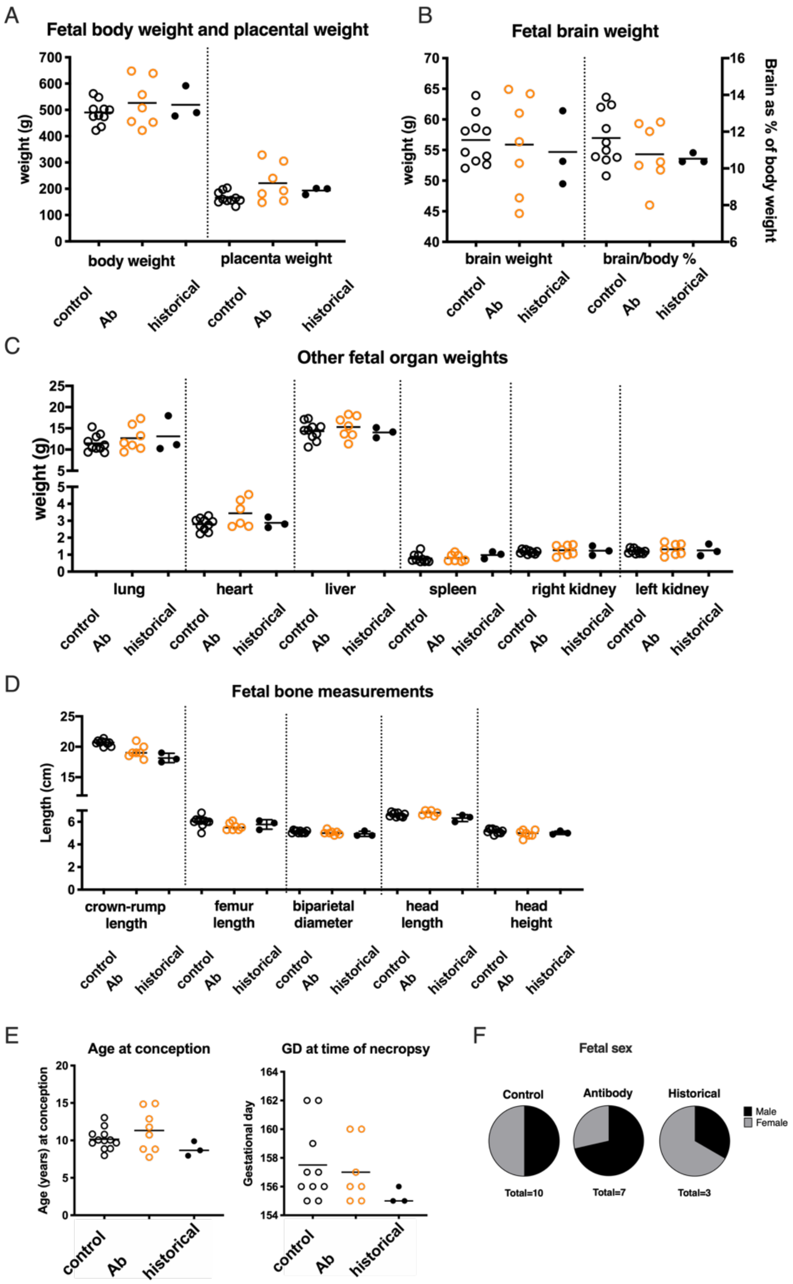


**Fig. S6. Fetal measurements at euthanasia from ZIKV infected macaques and from non-infected historical controls.** All measurements were collected on fetuses delivered via hysterotomy and immediately euthanized. The graphs include only fetuses that made it to the experimental endpoint (GD155-162) and exclude fetuses with early death. The historical controls are uninfected fetuses that also underwent intensive sampling as described earlier (11). **(A)** Fetal body and placental weights. **(B)** Fetal brain weights, including brain weight expressed as a percentage of total body weight. **(C)** Fetal organ weights. **(D)** Fetal bone measurements. **(E)** Age at conception and gestational day (GD) at time of euthanasia. **(F)** Sex of the fetuses.

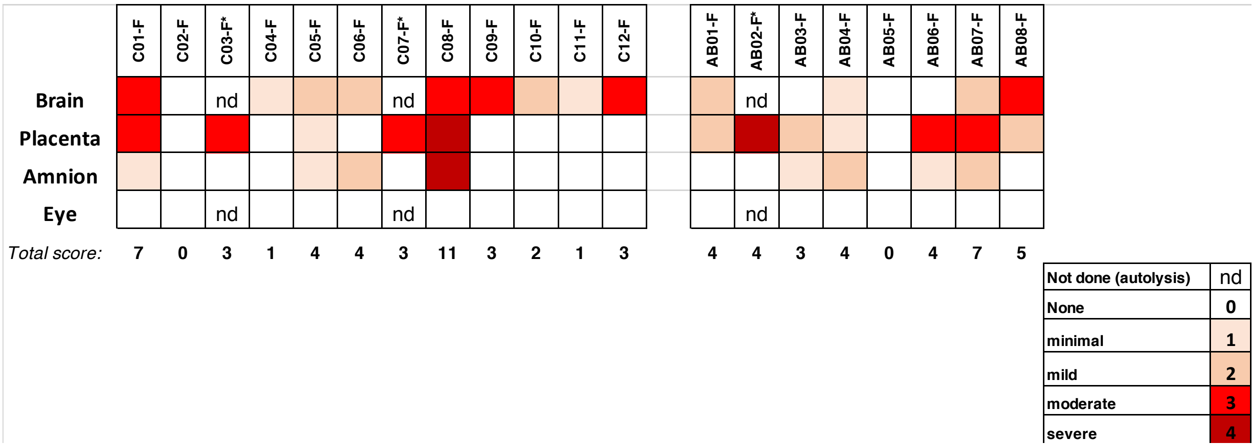


**Fig. S7. Fetal pathology scores.** Fetal brain (13-20 sections per animal), placenta (1-2 sections per animal), amnion (1-2 sections per animal) and eye (2 sections per animal) were evaluated blindly by a pathologist and were assigned a score of 0 to 4 based on severity (Tables S2, S3 and (21)). Asterisks indicate animals with early fetal death, for which some tissues could not be evaluated due to autolysis.

**Table S1. Histology of tissues from antibody treated dams.**

| Animal ID | Spleen | Mesenteric lymph node | Obturator lymph node | Inguinal lymph node | Uterus |
|---|---|---|---|---|---|
| AB01 | Moderate lymphoid hyperplasia | Mild lymphoid hyperplasia | Moderate hemorrhage in sinuses | Moderate lymphoid hyperplasia | Small numbers of perivascular lymphocytes in endometrium and myometrium (WNL) |
| AB02 | Mild lymphoid hyperplasia | Mild lymphoid hyperplasia | N/A | N/A | Small numbers of perivascular lymphocytes in the myometrium (WNL) |
| AB03 | Mild lymphoid hyperplasia | Severe lymphoid hyperplasia | WNL | Mild lymphoid hyperplasia | Small numbers of perivascular lymphocytes in endometrium and myometrium (WNL) |
| AB04 | Mild lymphoid hyperplasia | Mild lymphoid hyperplasia | Moderate macrophages in sinuses with hemosiderin | Moderate hemorrhage in sinuses; mild lymphoid hyperplasia | Thick decidua with hemosiderin-laden macrophages; mild to moderate numbers perivascular lymphocytes in endometrium and myometrium |
| AB05 | Moderate lymphoid hyperplasia | Mild lymphoid hyperplasia | Moderate hemorrhage in sinuses | Moderate hemorrhage in sinuses | Small numbers of perivascular lymphocytes in endometrium (WNL) |
| AB06 | Moderate lymphoid hyperplasia | Mild lymphoid hyperplasia | Moderate macrophages in sinuses | Mild lymphoid hyperplasia | Small numbers of perivascular lymphocytes in endometrium (WNL) |
| AB07 | Mild lymphoid hyperplasia | Moderate lymphoid hyperplasia | WNL | Mild lymphoid hyperplasia | Small numbers of perivascular lymphocytes in endometrium and myometrium (WNL) |
| AB08 | Mild lymphoid hyperplasia | WNL | WNL | WNL | Small numbers of perivascular lymphocytes in endometrium and myometrium (WNL) |

See in Van Rompay et al (ref. 21) for control pregnant macaques C01-C12; N/A is 'not available' and WNL is 'within normal limits'

**Table S2. Histology of fetal tissues from antibody treatment group.**

| Fetus ID | Central nervous system | Eye | Placenta | Amnion | Umbilical cord | Mesenteric lymph node | Lung | Liver | Kidney | Ileum |
|---|---|---|---|---|---|---|---|---|---|---|
| AB01-F | 19 sections: B5, B6, B7: segmental attenuation/loss of ependyma, mild, with gliosis | WNL | Basal plate and decidua have mild multifocal neutrophilic vasculitis and few interstitial neutrophils | WNL | WNL | WNL | Multifocal mild congestion and hemorrhage; rare squames | Mild EMH; mild glycogen; no GB | WNL | WNL |
| AB02-F | Severe autolysis | Severe autolysis | Placental separation; hemorrhage, fibrin and necrosis within decidua | WNL | N/A | N/A | Severe autolysis | Severe autolysis | N/A | N/A |
| AB03-F | 17 sections: B2: single focus mineral near ventricle | WNL | Basal plate and villous parenchyma multifocal areas of necrosis with neutrophils (slide 3-1, not the best sxn) | Very small numbers of neutrophils within the stroma | WNL | WNL | Rare squames | Rare EMH; moderate glycogen; no GB | WNL | WNL |
| AB04-F | 17 sections: B2, B3: rare individual cells in ventricular lumen; B4, B5: segmental attenuation/loss of ependyma, mild, with gliosis; cystic choroid plexus | WNL | Increased syncytial knots | Cystic change to epithelium; hemorrhage and hemosiderin-laden macrophages in stroma; thick decidua | WNL | WNL | Mild squames | Moderate EMH; moderate glycogen | WNL | WNL |
| AB05-F | 17 sections: B5: single focus mineral near ventricle | WNL | WNL | WNL | WNL | WNL | Rare squames | Mild EMH; moderate glycogen | WNL | WNL |
| AB06-F | 19 sections: B6: LGN less layering | WNL | Multifocal necrosis/loss of trophoblasts with neutrophils along fetal plate; large mid-disc infarct extending from the basal plate into the villous parenchyma with necrosis/mineralization of the basal plate and decidua; increased syncytial knots | Single area of moderate numbers of neutrophils within decidua | WNL | WNL | Mild squames | Rare EMH; moderate glycogen | WNL | WNL |
| AB07-F | 20 sections: Parietal lobe (10): multifocal mineralization near ventricle and adjacent neuronal rests; B2,B3: single rest mineralization | WNL | Severe decidual thickening with hemorrhage, edema and necrosis; multifocal ischemic necrosis of placenta full thickness (chronic and acute); mild lymphocytes and moderate neutrophils in decidua | Severe decidual thickening with multifocal small areas of hemorrhage and fibrin; mild lymphocytes | WNL | N/A | Moderate hemorrhage within alveoli | Mild EMH; moderate glycogen | WNL | Moderate hemorrhage within lumen; cytoplasmic protein droplets in enterocytes |
| AB08-F | 19 sections: Parietal lobe (10), lateral ventricle (15), B3, B4, B5: single to few focus/foci periventricular mineralization; Lat ventricle (15), B4: mild loss of ependyma with gliosis; B6, B7: moderate ependymal loss | WNL | Fetal plate multifocal trophoblast necrosis with fibrin and neutrophils, multifocal very mild neutrophils in fetal vessels; decidua multifocal necrosis with neutrophils | WNL | WNL | WNL | Moderate squames | Rare EMH; mild glycogen | WNL | WNL |

See in Van Rompay et al (ref. 21) for control pregnant macaques C01-C12; N/A is 'not available', WNL is 'within normal limits' and EMH is extra-medullary hematopoiesis

**Table S3. Fetal pathology scoring.**

| Score | Placenta/fetal membranes | Fetal neuropathology |
|---|---|---|
| **0** | none - no lesions observed | none - no lesions observed |
| **1** | minimal - focal, small lesion in section, not observed in control animals but significance questionable, could be background lesion | minimal - focal, small lesion in one brain section, not observed in control animals but significance questionable, could be background lesion |
| **2** | mild – mild, multifocal lesions in section, single layer | mild – mild ependymal loss in multiple brain sections +/- very few mineral foci |
| **3** | moderate – mild to moderate lesions extensively throughout section, multiple layers | moderate – mild to moderate ependymal loss in multiple sections, mineralization +/- cuffing |
| **4** | severe – widespread lesions throughout section, multiple layers | severe – widespread lesions in multiple brain sections |

**Table S4. Macaques used in the study.**

| Animal ID | Treatment | Date of birth | Sex | Total prior pregnancies | Delivery of live infants | Age at necropsy (in years) |
|---|---|---|---|---|---|---|
| **LOW DOSE SUBCUTANEOUS ZIKV CHALLENGE (NONPREGNANT)** | | | | | | |
| 6467 | Z004+Z021 GRLR | 5/2/15 | M | n.a. | n.a. | 2.8 |
| 6476 | Z004+Z021 GRLR | 5/11/15 | M | n.a. | n.a. | 2.8 |
| 6408 | Z004+Z021 GRLR | 3/7/15 | M | n.a. | n.a. | 3.3 |
| 6425 | Z004+Z021 GRLR | 3/30/15 | M | n.a. | n.a. | 3.2 |
| 6430 | Z004+Z021 GRLR | 4/3/15 | M | n.a. | n.a. | 3.2 |
| 6446 | Z004+Z021 GRLR | 4/17/15 | M | n.a. | n.a. | 3.2 |
| 6543 | Z004+Z021 GRLR/LS | 4/21/16 | F | n.a. | n.a. | 2.8 |
| 6575 | Z004+Z021 GRLR/LS | 7/2/16 | F | n.a. | n.a. | 2.6 |
| 6522 | Z004+Z021 GRLR/LS | 3/19/16 | F | n.a. | n.a. | 2.9 |
| 6525 | Z004+Z021 GRLR/LS | 4/1/16 | F | n.a. | n.a. | 2.8 |
| 6553 | Z004+Z021 GRLR/LS | 5/6/16 | F | n.a. | n.a. | 2.7 |
| 6642 | Z004+Z021 GRLR/LS | 4/27/17 | M | n.a. | n.a. | 1.8 |
| 6496 | untreated control | 6/11/15 | M | n.a. | n.a. | 3.0 |
| 6315 | untreated control | 4/7/14 | M | n.a. | n.a. | 4.0 |
| 6500 | untreated control | 6/18/15 | M | n.a. | n.a. | 2.8 |
| 6501 | untreated control | 6/21/15 | M | n.a. | n.a. | 3.0 |
| **LOW DOSE SUBCUTANEOUS ZIKV CHALLENGE (PREGNANT)** | | | | | | |
| AB01 | Z004+Z021 GRLR/LS | 3/19/04 | F | 10 | 9 | 15.3 |
| AB02 | Z004+Z021 GRLR/LS | 2/21/04 | F | 8 | 7 | 15.1 |
| AB03 | Z004+Z021 GRLR/LS | 4/28/08 | F | 6 | 5 | 11.1 |
| AB04 | Z004+Z021 GRLR/LS | 3/25/10 | F | 4 | 2 | 9.3 |
| AB05 | Z004+Z021 GRLR/LS | 3/15/06 | F | 6 | 3 | 13.3 |
| AB06 | Z004+Z021 GRLR/LS | 4/21/07 | F | 4 | 3 | 12.1 |
| AB07 | Z004+Z021 GRLR/LS | 4/3/10 | F | 3 | 0 (1 abortion, 2 experimental harvests) | 9.3 |
| AB08 | Z004+Z021 GRLR/LS | 5/1/11 | F | 2 | 2 | 8.2 |

See in Van Rompay et al. (ref. 21) for control pregnant macaques C01-C12; n.a. is 'not applicable'.

**Table S5. List of primers used in this study.**

| Primer ID | Primer sequence | Comments |
|---|---|---|
| Generation of pZIKV/HPF/CprM*PRVABC59E* | | |
| RU-O-24379 | ACTTGGTCATGATACTGCTGATTGCCCCGGCATACAGCAT CAGGTGCATAGGAGT | Forward with BspHI |
| RU-O-24380 | TTCGAACCGCGGCTGGGTCCTATTAAGCAGAGACAGCTG TGGATAAGAAGATC | Reverse with SacII |
| Mutagenesis of pZIKV/HPF/CprM*PRVABC59E* to produce mutant ZIKV RVPs | | |
| DFRp1477 | ACTTGGTCATGATACTGCTGATTG | Forward outer with BspHI |
| DFRp1584 | AACCGCGGCTGGGTCCTATTAAGCAGA | Reverse outer with SacII |
| DFRp1505 | GACAGTCACAGTGGAGTTACAGTACGCAGGGAC | Forward V330L |
| DFRp1506 | GTCCCTGCGTACTGTAACTCCACTGTGACTGTC | Reverse V330L |
| DFRp1503 | GTGTCATACTCCTTGTGTGCTGCAGCGTTCACA | Forward T309A |
| DFRp1504 | TGTGAACGCTGCAGCACACAAGGAGTATGACAC | Reverse T309A |
| DFRp1557 | GTGTCATACTCCTTGTGTATTGCAGCGTTCACA | Forward T309I |
| DFRp1558 | TGTGAACGCTGCAATACACAAGGAGTATGACAC | Reverse T309I |
| Mutagenesis of pZIKV/HPF/CprM*E* to produce mutant ZIKV RVPs | | |
| DFRp1477 | ACTTGGTCATGATACTGCTGATTG | Forward outer with BspHI |
| DFRp1481 | CGAACCGCGGGCCCTCTAGATCAA | Reverse outer with SacII |
| DFRp1546 | GCAGGGGCAGATGGACCTTGC | Forward T335A |
| DFRp1547 | GCAAGGTCCATCTGCCCCTGC | Reverse T335A |
| DFRp1548 | GCAGGGATAGATGGACCTTGC | Forward T335I |
| DFRp1549 | GCAAGGTCCATCTATCCCTGC | Reverse T335I |
